# Improved myelin clearance and cognitive outcomes after TBI in female mice are mediated by ovarian steroids and sex chromosomes

**DOI:** 10.64898/2026.01.28.702273

**Authors:** Daniel Pinto-Benito, Carmen Paradela-Leal, Nuria Cano-Adamuz, Daniela Grassi, Iñigo Azcoitia, Sofia Grade, Maria-Angeles Arevalo

## Abstract

Traumatic brain injury (TBI) causes sex-specific memory deficits, yet the underlying mechanisms are not fully understood. Using a mouse TBI model, we investigated the role of reactive astrocytes in sex-specific outcome. TBI provoked long-term contextual memory impairment in males and ovariectomized females, but not in intact females. The synthetic steroid tibolone preserved memory and cFos+ neuronal density in the hippocampus of ovariectomized females. Hormone deprivation upregulated astrocytic GFAP and S100B, reduced Homer1, and impaired myelin phagocytosis by astrocytes in females. These effects were counteracted by tibolone. In Four-Core-Genotype mice, memory loss correlated with reduced astrocytic myelin uptake and neuronal activity in XX males and XY female animals. Astrocyte transplantation showed that female astrocytes exhibit superior myelin clearance capacity, especially in female brain environments, though they outperform male astrocytes in both sex contexts. These findings identify astrocyte-mediated myelin phagocytosis as a key mechanism for memory preservation after TBI, governed by both hormonal and chromosomal sex factors.

## Introduction

Traumatic brain injury (TBI) is a leading cause of death and long-term disability worldwide and significantly increases the risk of cognitive decline and neurodegenerative diseases [1]. Both clinical and preclinical studies consistently report sex differences in TBI outcomes, with females, on average, exhibiting greater resilience and improved recovery compared to males [2]. This protective advantage is evident in both cellular pathophysiology and functional outcomes, yet the biological basis of the sex-based disparities remains poorly understood. Gonadal hormones, particularly estrogens and progesterone, have long been implicated in neuroprotection following brain injury [3]. These hormones can modulate gene expression, synaptic plasticity, and inflammatory responses in the brain. Notably, both neurons and glial cells, including astrocytes and microglia, express steroid hormone receptors, enabling direct hormonal modulation [4, 5]. Synthetic steroidal compounds such as tibolone have been developed to mirror the beneficial effects of natural hormones while minimizing off-target effects. Tibolone, which acts through estrogen, progesterone, and androgen receptors [6], has shown great promise in preclinical studies wherein it reversed dendritic spine loss, reduced oxidative stress, and attenuated neuroinflammation after TBI [7-9]. Given the well-documented contribution of neuroinflammation to cognitive impairment [10], investigating how tibolone modulates such impaired cognitive functions and glial responses may yield new therapeutic insights.

One key function of glial cells after TBI is the clearance of cellular debris, particularly myelin fragments. Myelin debris acts as a potent immune signal, activating a complex repertoire of receptors and downstream pathways [11]. Accumulation of myelin debris at the lesion site is thus detrimental, as it amplifies inflammation, impairs remyelination, and disrupts tissue homeostasis. Effective clearance of such debris has been linked to improved hippocampus-dependent memory, particularly in aging brains [12, 13]. While microglia are classically regarded as the primary phagocytes of the CNS, a growing body of evidence indicates that astrocytes also contribute to phagocytosis under both physiological and pathological conditions. Both our prior in vitro work and in vivo studies demonstrate that astrocytes can internalize and degrade myelin debris, and importantly, that their phagocytic capacity is determined by sex [14-16]. Astrocytes are increasingly recognized as central players in both injury response and repair. Despite the emerging evidence, the role of astrocytes in mediating sex-specific outcomes after TBI remains largely unexplored. Addressing this gap is critical, as astrocytes may represent a novel cellular target for sex-informed therapeutic strategies.

In this study, we set out to investigate how sex-specific factors – hormonal and chromosomal – influence astrocyte function and cognitive resilience after TBI. To do so, we employed a multifaceted experimental approach: 1) gonadectomy and Tibolone treatment in wildtype mice to examine the role of gonadal hormones in preserving memory and modulating astrocyte activity after TBI; 2) the Four-Core-Genotype (FCG) mouse model to disentangle the contributions of sex chromosomes versus gonadal hormones on astrocyte reactivity, phagocytosis, and on memory test performance; 3) crossed-sex astrocyte transplantation, using fluorescently labeled astrocytes from male and female Aldh1l1-EGFP mice, to evaluate the impact of sex-specific brain environmental signals in shaping astrocytic phagocytosis and myelin clearance.

Our findings reveal a novel astrocyte-centered mechanism that contributes to sex differences in recovery after brain trauma. Specifically, we show that female astrocytes display enhanced myelin phagocytosis, particularly within female brain environments, which is associated with preserved neuronal activity and contextual memory. These results uncover an underappreciated dimension of astrocyte function in TBI and suggest that modulating astrocytic phagocytosis may offer a promising therapeutic strategy to improve cognitive outcomes in a sex-specific manner.

## Materials and Methods

### Animals

14-week-old C57-BL/6J, Four Core Genotype (FCG, a kind gift from A.P. Arnold, UCLA, USA), and Aldh1l1-EGFP (RRID: MMRRC_011015-UCD) mice raised in our in-house colony at the Cajal Institute were used for this study. In FCG mice, males (XYm) have the testis-determining gene Sry deleted from the Y chromosome, and inserted into an autosome [17]. XYm mice were bred to C57BL/6J WT females to produce FCG mice of four genotypes: XYm/XXm (mice with Sry, gonadal males but with different chromosome combination) and XYf/XXf (mice lacking Sry, thus gonadal females but with different sex chromosome combination) [18]. Aldh1/1-GFP transgenic mice express GFP under the Aldh1l1 promoter, providing a way to specifically label and study astrocytes. Animals were housed under controlled temperature (22 ± 2 °C) and light conditions (12-h light/dark cycle) and with food and water available *ad libitum*. All the procedures applied to the animals used in this study followed the European Parliament and Council Directive (2010/63/EU) and the Spanish regulation (R.D. 53/2013 and Ley 6/2013, 11th June) on the protection of animals for experimental use and were approved by our institutional animal use and care committee (Comité de Ética de Experimentación Animal del Instituto Cajal) and by the Consejería del Medio Ambiente y Territorio (Comunidad de Madrid, PROEX 059.5/21). Genotyping of FCG mice was performed by RT-PCR detection of Sry and Ssty (located in the Y chromosome) gene transcripts.

Male and female mice were randomly assigned to four experimental groups: Sham: mice without injury; Injured: mice that received a TBI; GNX+Inj mice that were gonadectomized prior to the injury; GNX+Inj+Tib: mice that after being gonadectomized and injured, were treated with Tibolone. The experimental design is represented in Fig. 1A.

**Fig. 1:**
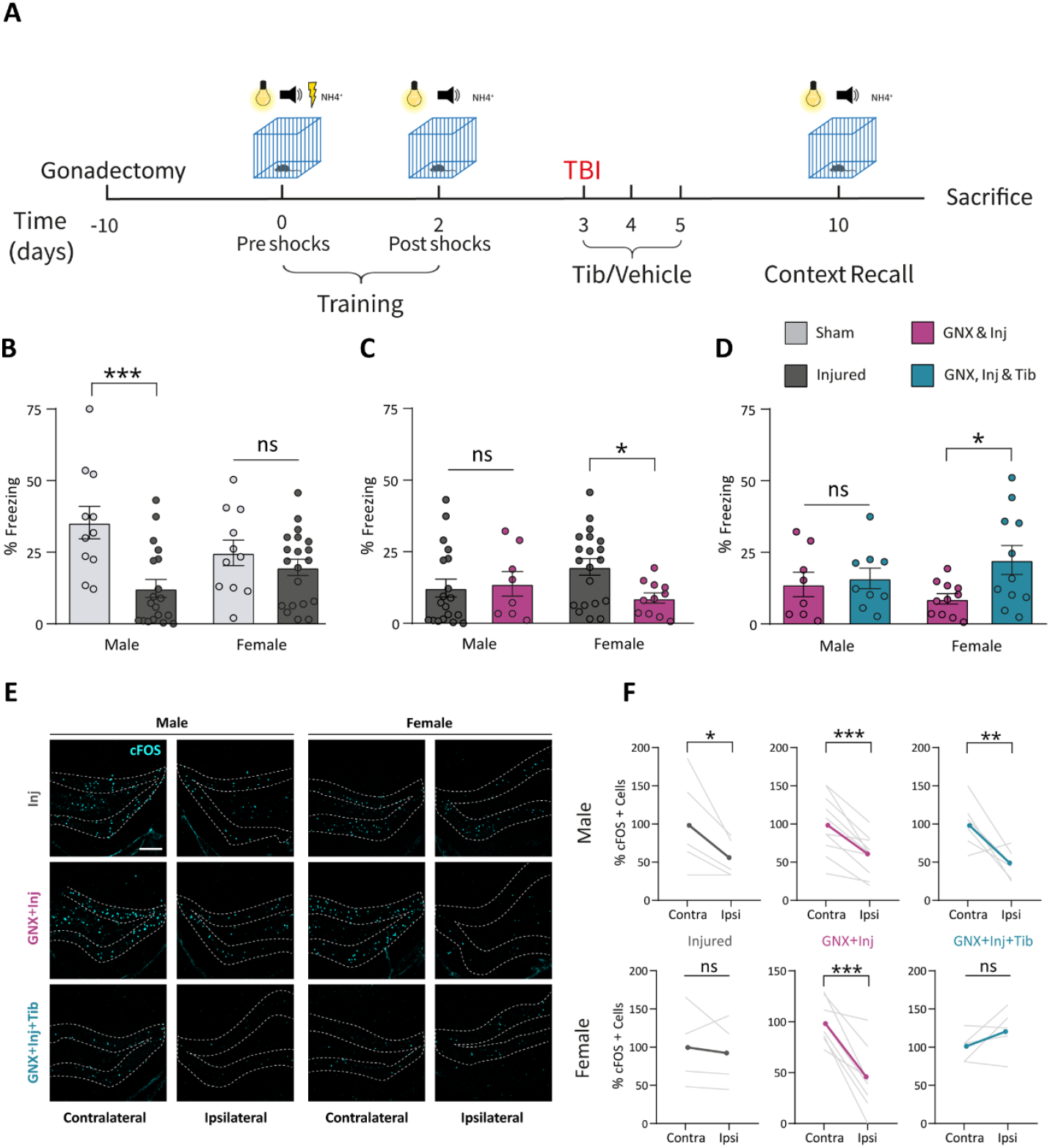
Traumatic brain injury affects hippocampal memory in a manner influenced by sex and steroid hormones. (**A**) Time curse of the experimental procedure. Some animals were castrated at least 10 days before starting the protocol (gonadectomy, GMX). All animals received habituation to the experimental room and experimenter during this period. Freezing behavior was quantified in a training phase, before (Pre-shocks) and after (Post-shocks) being submitted to electric shocks. 24h later, a controlled cortical impact (TBI) was made over the medial cortex. Vehicle or tibolone (tib) were administered at the time of injury and 24h and 48h post-injury. On day 7 post-TBI, animals were evaluated for memory recall and sacrificed. Effect of TBI (Inj) (**B**), gonadectomy (**C**), and tibolone treatment (**D**) on freezing behavior (% of time), respectively. (**E**) Representative images of neuronal activation in the hippocampus immunostained with anti-cFos antibody (cyan). (**F**) Quantification of the cFos+ cells in the contralateral and ipsilateral hippocampus of the evaluated groups. Note the reduction in all male groups and castrated females. Data in B, C, and D are presented as mean ± SEM; two-way ANOVA followed by Fisher’s multiple comparisons post hoc test, n ≥ 8 animals per group. Data in F are presented as individual values; paired t-test. n = Individual values for each histological section (equivalent histological sections from a minimum of 4 animals). *p < 0.05; ***p < 0.001; ns = non-significant. Scale bar 200µm.

### Gonadectomy

Animals were castrated under isoflurane anesthesia at least 10 days before behavioral tests in order to ensure total clearance of circulating gonadal hormones.

For female ovariectomy, a bilateral 0,5 cm incision was performed in the skin in the lower back region of the animals. Ovaries were recognized through the posterior wall of the abdominal cavity and removed. To avoid hemorrhages and infections, the union of the ovary to the oviduct was cauterized (Alsatom, Alsa), and the skin was sutured with absorbable suture thread.

For male orchidectomy, bilateral 0,5 cm incisions were made on the ventral side of the scrotum. Testicles were identified through the perineal floor and removed by excision. The union of the testicle to the ductus deferens was cauterized to prevent bleeding and infection, and the skin of the scrotum was sutured with absorbable suture thread.

### Traumatic brain injury

Mice were anesthetized with isoflurane (induction at 5% and maintenance at 2% in O_2_) and placed in a stereotaxic apparatus (David Kopf Instruments) at 37°C. A sagittal incision of about 1.5 cm was made along the midline of the animal’s dorsal skull surface. After cleaning the exposed skull, the head was leveled and the intended injury site (somatosensory cortex) was marked at the following coordinates on the Paxinos and Franklin mouse brain atlas (Bregma -1.5 cm antero-posterior, -1.5 cm medio-lateral) on the left hemisphere (Supplementary Fig. 1A). Using a 2.5 mm drill, a craniotomy was performed, taking care to not damage the meninges. After cleaning the area, a 2.3 mm hammer connected to the injury system (Impact OneTM, MyNeuroLab, Leica) was carefully lowered until contacting with the meninges. The firing pin was then retracted and the piston was lowered 1mm. Afterward, the controlled cortical impact was performed with a depth of 1 mm on the desired coordinates, at a speed = 3.6 m/s and with a latency of 0.1 s. Next, the hemorrhage was cleaned and the head skin was sutured with 5/0 surgical suture. In sham animals, the cranial surface was drilled, but no injury was performed.

### Tibolone treatment

Animals received subcutaneous injections of tibolone (10006321, Cayman Chemical, 0.04 mg/Kg) or vehicle (corn oil) immediately, 24 h and 48 h after brain injury (Fig. 1A). The dose and protocol of tibolone administration were selected based on our previous studies [8] and on the posology used in clinical practice (2.5 mg/day).

### Fear conditioning test

For 3 consecutive days prior to the start of the Fear Conditioning protocol, mice were placed into behavior testing room and handled by the experimenter as a process of habituation to the experimental conditions. At day 0, animals were placed in the fear conditioning chamber (Ugo Basile), with barred floor and context cues such as visual cues (high-intensity light and semi-transparent walls), sound cues (white noise), and smell cues (1% NH_4_^+^) with the aim of making the context easily recognizable by mice. Freezing behavior was monitored using Anymaze software (Stoelting). The training phase consisted of placing the animals for 3 minutes inside of the fear conditioning chamber until they received 3 rounds of electric shocks 0.3 mA/2 s. Later, animals were returned to their cage and housed for 2 days to consolidate the fear memory. Between animal tests, the behavioral cage was cleaned with 0.03% acetic acid in order to avoid unwanted odors that may influence animals’ behaviors.

After 2 days, animals were placed into the fear conditioning cage for 5 minutes without receiving shocks and the freezing response was analyzed using a freezing score scale. The score represents a measure of hippocampal-dependent contextual memory. Afterwards, mice were submitted to the TBI procedure, and 7 days post-injury, corresponding to the peak of astrogliosis [19], a final test was performed: mice were placed again into the behavioral cage for 5 minutes without receiving shocks, and the freezing behavior was analyzed. 24 h after finishing the test, animals were sacrificed for immunohistochemistry (IHC) or molecular biology studies.

### Immunohistochemistry

Mice were deeply anesthetized with pentobarbital (50 mg/kg body weight) and perfused through the left cardiac ventricle, first with pre-warmed (37 °C) 0.9% NaCl and then with 4% paraformaldehyde diluted in 0.1 M phosphate buffer (pH 7.4) for 5 min. Brains were post-fixed in the same fixative for 4 h at 4 °C and washed three times with 0.1 M phosphate buffer, pH 7.4 at room temperature. Coronal sections, 50 μm thick, were obtained using a Vibratome (VT 1000S, Leica Microsystems, Wetzlar, Germany).

IHC was carried out in free-floating sections under moderate shaking. All washes and incubations were done in 0.1 M phosphate buffer pH 7.4, containing 0.3% bovine serum albumin and 0.3% Triton X-100. After several washes, sections were incubated overnight at 4 °C with the following primary antibodies: mouse anti-GFAP-Cy3 (1:1000; Sigma, USA), rabbit anti-S100B (1:500; Abcam, UK), rabbit anti-Homer1 (1:500; Synaptic Systems, Germany), rabbit anti-Megf10 (1:500; Millipore, USA), rabbit anti-dMBP (1:500; Millipore, USA), mouse anti-NeuN (1:500; Millipore, USA), rabbit anti-cFOS (1:500; Abcam, UK). Sections were then rinsed in buffer and incubated for 2 h at room temperature with a goat anti-rabbit Alexa secondary antibody (1:1000; 488 nm or 594nm) and goat anti-mouse Alexa secondary antibody (1:1000; 488 nm or 594nm). After wash, brain sections were mounted on slides with Gerbatol antifade mounting medium with DAPI.

### Quantitative real-time polymerase chain reaction (qRT-PCR)

Following TBI and behavioral testing, cortices of both hemispheres were homogenized in TRIzol reagent (Invitrogen, Carlsbad, CA, USA), and the RNA was extracted. First-strand cDNA was prepared from 0.75 μg RNA using M-MLV reverse transcriptase (Promega, Alcobendas, Madrid) according to the manufacturer’s protocol. After reverse transcription, cDNAs were amplified by real-time PCR in 10 μl reaction volume using SYBR Green Master Mix (Applied Biosystems, Foster City, CA) and the ABI Prism 7500 Sequence Detection System (Applied Biosystems) with conventional Applied Biosystems cycling parameters (40 cycles of changing temperatures, first at 95 °C for 15 s and then 60 °C for 1 minute). All the primer sequences were designed using Primer Express software (Applied Biosystems) and are shown in Table 1. Primers were verified to amplify with a 95–100% efficiency by performing 4-point calibration curves and for each primer pair, an appropriate dilution of cDNA was chosen in order to achieve the same amplification efficiency as that of the housekeeping gene (Rpl13A). Changes in mRNA expression were calculated following the ΔCt method. mRNA levels were measured in both hemispheres, and mRNA expression was presented by subtracting the mRNA levels of the contralateral hemisphere from the ipsilateral one.

**Table 1.**
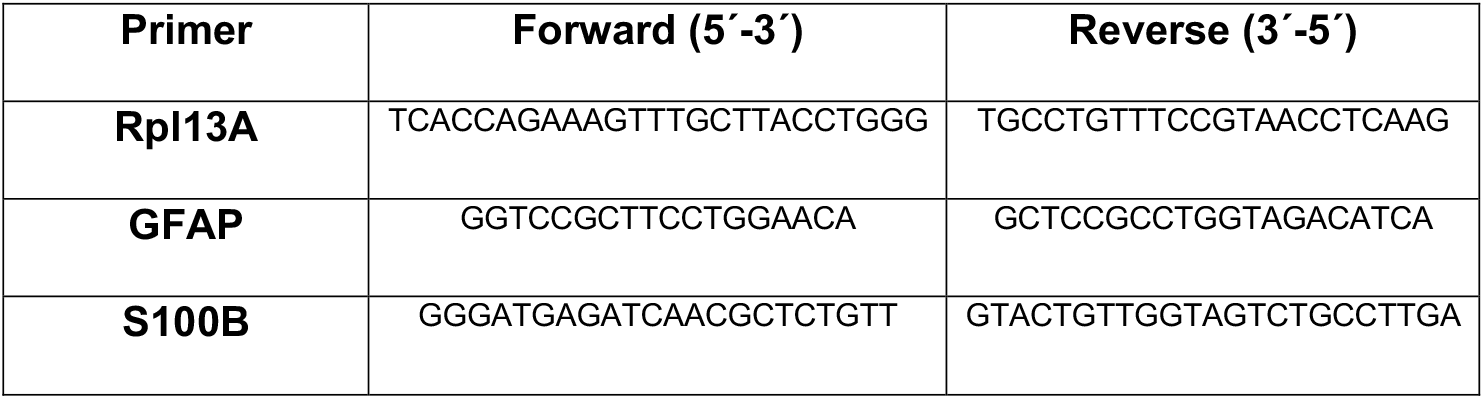
List of primers used in this study.

### Image acquisition and analysis

#### Image Acquisition

Confocal microphotographs were acquired as follows:

For whole cortex images, a Leica TCS-SP5 confocal system with a 20X objective was utilized. Images were captured in serial planes with a 3.5 μm-interval and with a resolution of 512×512 pixels (8-bit images). LASX software was employed to stitch together these images using the Tile Scan function.

To assess protein expression and phagocytosis, at least 3 images from each of 4 different animals per group, were acquired by an experimenter who was unaware of the experimental groups. The images were captured with the same acquisition settings and were taken at the same cortical region relative to the core of the injury (or corresponding cortical region in sham brains), of equivalent histological sections across animals. A Leica TCS-SP5 confocal system with a 63X objective was used and images were acquired in serial planes with a 0.5 μm-interval and with a resolution of 512×512 pixels (8-bit images).

#### Quantification of protein expression and phagocytosis

The intracellular location within astrocytes of proteins and debris was confirmed by a 3D reconstruction using IMARIS software (Bitplane, UK). Quantification of protein expression and phagocytosis was performed applying a threshold to systematically identify the entire GFAP area using Fiji imaging processing package (ImageJ 1.52n, National Institutes of Health, USA). The GFAP-positive area was used as ROI and an automatic macro with a smooth filter to discard background noise was applied to measure the integrated density (product of the fluorescence intensity of the corresponding protein by the area) of the protein of interest or the phagocyted particles inside the astrocytes. The obtained data were normalized to the values of the male injured group for comparative analysis.

For cFos evaluation, a projection of the complete confocal Z-stacks of the mouse hippocampus was performed and NeuN and cFos double-positive cells were counted in both hippocampal hemispheres.

### Astrocyte transplantation and quantification

For astrocyte transplantation, primary cultures of cortical astrocyte from male and female ALDH1/1-EGFP mice were prepared following the procedure described in [16]. These mice were generously provided by Gertrudis Perea’s laboratory and constitute a valuable tool for transplantation since astrocytes express the fluorescent reporter GFP endogenously.

Once the astrocyte cultures reached confluence, the cells were detached with 0.5% trypsin (Sigma-Aldrich), centrifuged, resuspended in DMEM at a concentration of 100.000 cells/μl and kept on ice for posterior injection. Cell transplantation followed a surgical procedure similar to the one described for TBI to expose the brain surface. A Hamilton syringe (point style = 3, Gauge = 33 and 30 mm needle length, Hamilton EEUU) was loaded with approximately 3μl of cell suspension without introducing bubbles. Four pulses of ∼30.000 cells each were injected at depths of 1 mm, 0.75 mm, 0.5 mm, and 0.25 mm from the brain surface adjacent to the area where the TBI later impacts i.e. at -1.5 mm AP, -2.7 mm ML relative to bregma (total 120.000 cells). After transplanting the cells, the cranial piece was placed back and the skin was sutured. Astrocyte transplantation was performed on mice of the same and of opposite sexes. Cell transplantation was performed 7 days before TB.

To quantify engulfment of myelin debris by transplanted astrocytes, cell volumes were reconstructed using the surface tool of IMARIS. A new channel was created to mask the debris inside those cell volumes. A 3D reconstruction of the myelin debris inside each cell was then made, and the value of their volume was obtained. Cells from at least 3 different animals were analyzed.

### Statistical analysis

All data shown in the figures are presented as the mean ± standard error of the mean (SEM). The size of the experimental groups corresponds to the number of animals, and it is indicated in each figure legend. Statistical analyses were carried out using GraphPad Prism software version 8.0 for Windows. The normality of the data was assessed with the Shapiro-Wilk test, to satisfy the assumption of normality. Data were evaluated by one- or two-way analysis of variance (ANOVA) with treatment and/or sex as independent variables. In two-way ANOVAs, the statistical significance of the effects of each independent variable and their interactions was tested. For those variables/interactions for which ANOVA p values were statistically significant, post hoc comparisons were performed with Fisher’s multiple comparisons test (otherwise, indicated in the figure legends). The statistical significance level was set at *p* < 0.05 in all cases.

## Results

### Traumatic brain injury causes a sex-dependent hippocampal memory impairment that is modulated by neuroactive steroids

To assess whether TBI leads to behaviorally relevant consequences, we tested for an effect on contextual fear memory as scheduled in Fig. 1A. Training for this memory test requires spatial exploration of a context and then delivery of an aversive stimulus, so animals learn to associate the contextual cues with an aversive stimulus, via formation of, so called, contextual fear memory [20]. No freezing behavior was detected before the exposure to the aversive stimulus (Supplementary Fig. 1A). As expected, during recall driven exclusively by contextual cues, male and female sham mice showed freezing responses after the training and, as previously described [21-23], females showed reduced freezing response compared with males (Supplementary Fig. 1B). Gonadectomy alone had no effect in the freezing response, neither in males nor in females (Fig. 1A (no TBI; no tibolone); Supplementary Fig. 1B). In animals subjected to TBI (Supplementary Fig. 1C), the behavioral test performed 7 dpi, indicated that the injury provoked a significant reduction in freezing levels in males while it had no effect in females (Two-way-ANOVA, F(interaction)=5.287, p=0.0252; followed by Fisher’s multiple comparisons test; sham males vs. injured males p=0.0001) (Fig. 1B and Supplementary Fig. 1D). Next, we evaluated the effect of gonadectomy on the memory of the injured animals (GNX+Inj). The results indicated that castration did not affect the freezing response of males. In contrast, gonadectomy produced a reduction in freezing levels in females (Unpaired t test, p= 0,0064) (Fig. 1C). Notably, this effect was counteracted by tibolone treatment while in males, hormone replacement did not affect the freezing rate (Two-way-ANOVA, F(treatment)=4.288, p=0,0458; followed by Fisher’s multiple comparisons tests; GNX+Inj+Tib vs. GNX+Inj females, p=0.0269) (Fig.1D and Supplementary Fig. 1E and 1F). These results demonstrated that, regardless of the presence or absence of steroid hormones, TBI consistently causes long-term memory impairment in male animals, while in females an impairment is achieved only in castrated individuals pointing to a protective role of gonadal hormones in females that enable preservation of hippocampal memory.

### Sex differences in hippocampal activity during memory recall in mice subjected to TBI

We next tested whether steroid hormones affected neuronal activity induced by reactivation of the contextual fear memory, in the hippocampal dentate gyrus, a structure laying just underneath the cortical injury (Supplementary Fig. 1A), and wherein memory traces for contextual fear were previously identified [24-26]. For this purpose, we measured the number of cFos+ neurons in the dentate gyrus, as the immediate early gene cFos is associated with neuronal activity (Supplementary Fig. 1G). In line with our behavioral results, TBI caused in males a drastic decrease in the number of cFos+ neurons in the ipsilateral hemisphere compared to the contralateral one (paired t-test, p=0.0261), while no significant differences were observed in females (Fig. 1E and 1F). Regardless of whether males have been only inflicted with TBI (Inj) or gonadectomized and injured (GNX+Inj) or gonadectomized, injured and treated with tibolone (GNX+Inj+Tib), the ipsilateral hippocampus shows lower levels of cFos+ neurons compared to the contralateral (paired t-test, p=0.0006 for GNX group, p=0.0240 for Tib group). Surprisingly, injured females with intact ovaries showed similar levels of active neurons between the two hippocampi. However, the injury effectively reduced the number of active neurons in ovariectomized females (paired t-test, p= 0.0001) and tibolone treatment equalized the levels in both hemispheres (paired t-test, p=0.1087) (Fig. 1F).

Altogether, these findings suggest that steroid hormones produced in the female gonads prevent the hippocampal memory loss associated with brain injury, by maintaining functional neuronal circuits that underlie memory retention.

### Male and female hormonal status influences astrocyte reactivity in TBI

To assess the possible role of astrocytes underlying female-specific neuroprotection after TBI, we evaluated at 7 dpi, the reactivity of astrocytes surrounding the injury. We had previously shown that intact female and male mice have similar densities of GFAP immunoreactive cells at given distances to a stab wound injury [27]. Here, we analyzed astrogliosis in gonadectomized mice subjected to TBI and the effect of tibolone treatment. We assessed the mRNA expression level of GFAP and of S100B, indicative of astrocyte reactivity, in the tissue around the lesion (Fig. 2A-D). Two-way ANOVA did not reveal any significant difference in the GFAP mRNA expression between sexes. Nevertheless, tibolone significantly reduced the increased GFAP levels in gonadectomized females (One-way ANOVA, F(treatment)=15.72, p=0.0012; followed by Fisher’s multiple comparisons test, p= 0.0126, GNX+Inj vs. Injured and p=0.0010, GNX+Inj+Tib vs. GNX+Inj) (Fig. 2B). In addition, we found that S100B mRNA expression was significantly higher in injured males compared to injured females and gonadectomy reduced S100B levels only in males (Two-way ANOVA, F(sex)=7.956, p=0.0097; followed by Fisher post-hoc test (p=0.0221, Inj vs. GNX+Inj males). Tibolone treatment had no effect on either sex (Fig. 2D).

**Fig. 2:**
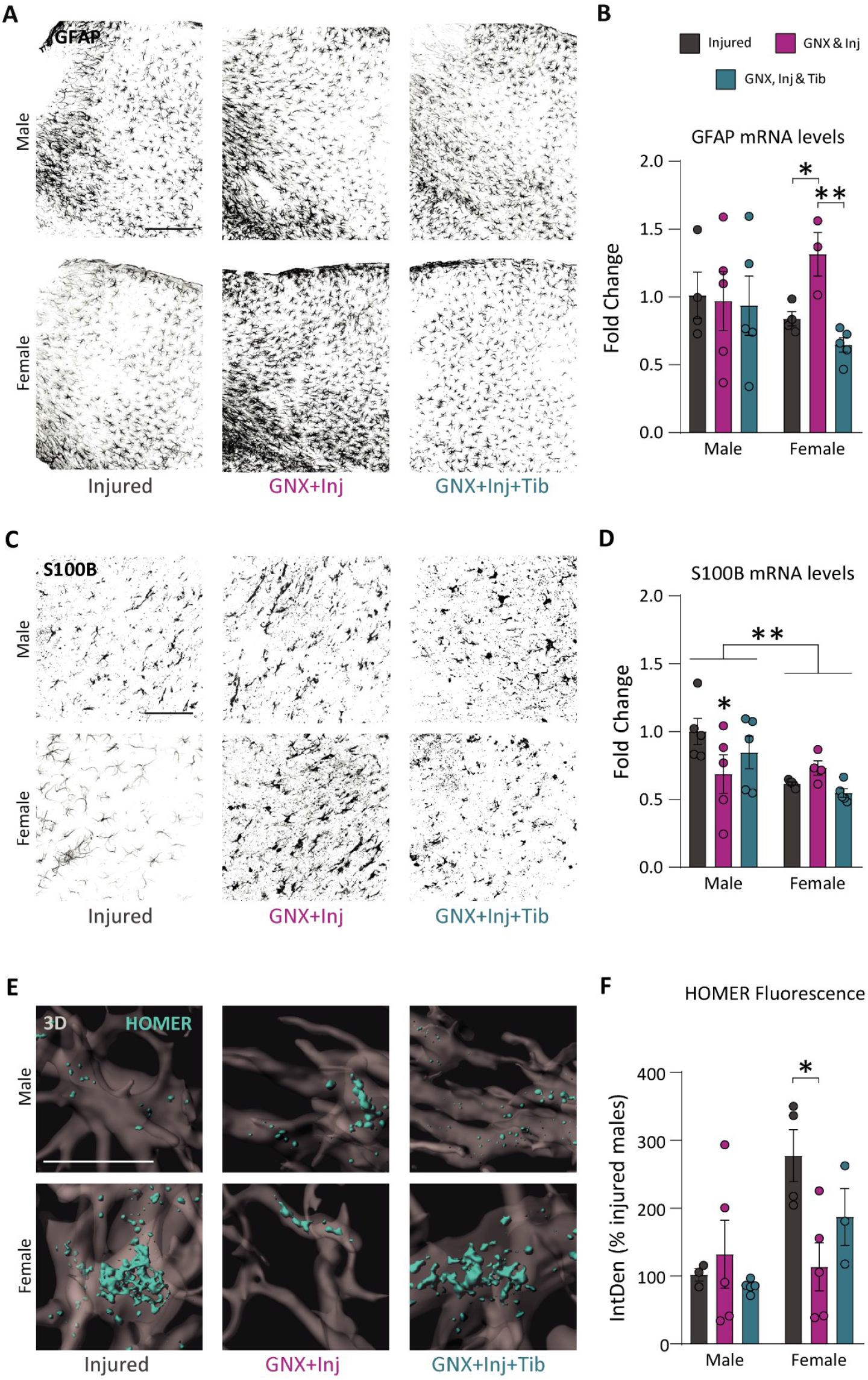
Astrogliosis and astrocytic Homer1 expression in response to injury are hormone and sex-dependent. (**A, C**) Representative confocal images of GFAP (black) and S100B (black) IHC signal in the cortex. (**B, D**) Quantification of GFAP and S100B mRNA levels in the tissue lysates, measured by qRT-PCR. Data normalized to housekeeping gen and presented as fold change relative to the Inj male group. (**E**) IMARIS 3D reconstructions of Homer1 (cyan) expression in astrocytes (gray). (**F**) Quantification of the percentage of HOMER integrated density (IntDen) inside the GFAP+ cells. Data are presented as the mean ± SEM; two-way ANOVA followed by Fisher’s multiple comparisons post hoc test. n ≥ 3 animals per group. *p < 0.05; **p < 0.01; ***p < 0.001. Scale bars: 250µm in (A), 100µm in (C) and 15µm in (E).

Homer1 is expressed in neurons and is a major component of synapses, however recent evidence also demonstrates Homer1 expression in astrocytes. The expression of Homer1 in astrocytes has been associated with neuroprotection by promoting non-inflammatory phenotypes in astrocytes [28]. Hence, examined the expression of Homer1 in astrocytes of mice subjected to TBI, by IHC (Supplementary Fig. 2). We measured the integrated density of Homer1 within GFAP+ cells surrounding the injury (Fig. 2E and F). Two-way ANOVA showed an interaction effect between sex and treatment (F=3.557, p=0.0451). Multiple comparison analysis using Fisher’s post hoc test revealed that Homer1 expression is not affected by the castration or tibolone treatment in male astrocytes. However, ovariectomy of females significantly reduces Homer1 levels compared to non-castrated animals (p=0.0203) with tibolone partially preventing this downregulation (Fig. 2E and F).

Together these results indicate that although TBI causes an increase in astrocyte reactivity in both sexes, the presence of ovarian or supplemented steroid hormones avoids excessive astrogliosis in female mice.

### Phagocytosis by reactive astrocytes is influenced by sex hormones

Considering the results described above, which correlated sex hormones with astrocytic reactivity, expression of Homer1, neuronal activity and contextual memory, we asked whether they also modulate astrocyte phagocytic capacity, a process that supports a healthier environment. To address this, we assessed myelin phagocytosis by astrocytes near the lesion, by quantifying dMBP fluorescence within GFAP+ cells, using IHC (Supplementary Fig. 3A). Two-way ANOVA revealed a significant interaction effect between sex and treatment (F=4.310, p=0.0252) and an effect of the treatment (F=3.507, p=0.0461). While no differences were observed among male groups, Tukey’s multiple comparisons test revealed that castration of females significantly reduced myelin uptake by astrocytes (p=0.0076), being this effect abrogated by tibolone treatment (Fig. 3A and 3B).

**Fig. 3:**
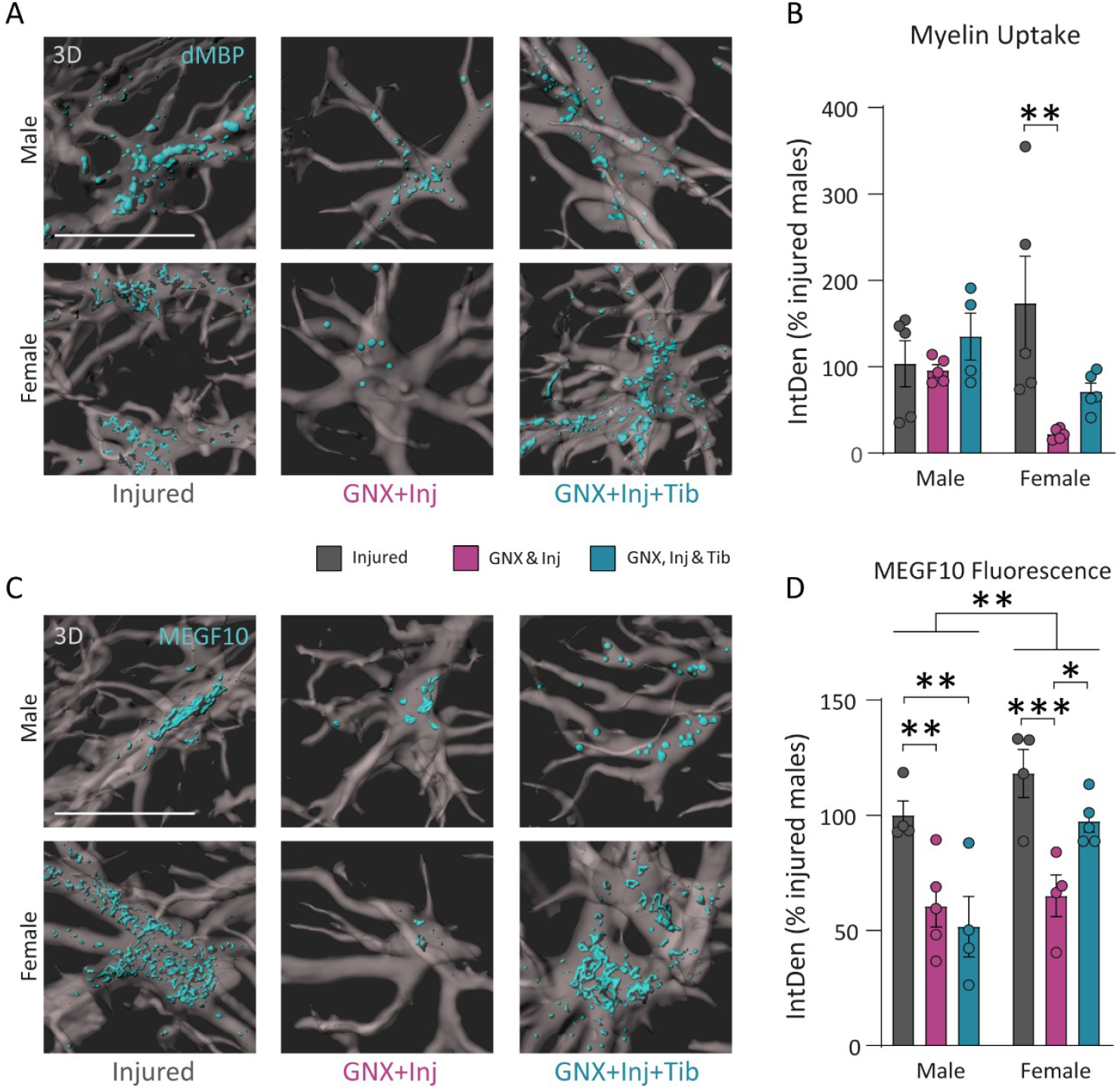
Myelin uptake and phagocytic receptor (Megf10) expression are conditioned by sex and gonadal hormones. (**A, C**) IMARIS 3D reconstructions of dMBP and MEGF10 (cyan) immunoreactivity within GFAP+ astrocytes (gray). (**B, D**) Quantification of engulfed myelin and Megf10 protein expression was evaluated by measuring dMBP and Megf10 fluorescence intensity (IntDen) wihin the astrocytes. Data are presented as mean ± SEM; two-way ANOVA followed by Fisher’s multiple comparisons post hoc test. n ≥ 4 animals per group. *p < 0.05; **p < 0.01; ***p < 0.001. Scale bars 15µm.

To further investigate how steroid hormones regulate astrocyte phagocytosis, we measured Megf10 fluorescence intensity in GFAP+ cells in the perilesional tissue by IHC (Fig. 3C, 3D and Supplementary Fig. 3B). Megf10 is a transmembrane receptor in astrocytes that recognizes “eat-me” signals and initiates internalization of targets and engulfment. Two-way ANOVA followed by Fisher’s multiple comparisons test revealed effects of sex (F=9.763, p=0.0053) and treatment (F=14.07, p=0.0002) without interaction, on the levels of Megf10. Gonadectomy significantly reduced Megf10 levels in astrocytes near the lesion in both sexes (p=0.0047, Injured males vs. GNX+Inj males and p=0.0006, Injured females versus GNX+Inj females). Tibolone treatment significantly reversed this reduction only in females (p=0.0171, GNX+Inj females vs. GNX+Inj+Tib females) while it had no effect in males (Fig. 3D).

Together these results suggest a crucial role of steroid hormones in astrocyte phagocytosis, mainly in females. In the presence of these hormones (produced in the gonads or administered as treatment) female astrocytes exhibit enhanced myelin phagocytosis. This likely creates a less harmful microenvironment, potentially preventing neuronal death and promoting neuronal survival.

### Chromosomal complement contributes to sex differences in memory, hippocampal neuron activity, astrocyte reactivity, and phagocytosis in TBI

To decipher the contribution of gonadal hormones and sex chromosomes in the outcome of TBI, we performed experiments using the FCG model (Supplementary Fig. 4A). FCG animals were submitted to the experimental paradigm described in Fig. 4A. As expected, animals did not exhibit freezing levels before exposure to the aversive stimulus (Supplementary Fig. 4B). Nevertheless, electric shocks caused a significant increase in freezing levels, with XX animals exhibiting reduced freezing time compared to the XY counterparts (Supplementary Fig. 4C). After TBI, Two-way ANOVA followed by Fisher’s multiple comparisons test revealed an interaction between chromosomes and gonads (F=19.17, p<0,000) with XYf and XXm showing a significant reduction in freezing compared to chromosomal counterparts (p=0.0311 and p=0.0002, respectively). The analysis by Mann Whitney test also detected a significant difference between male groups (p=0.0061) and between female groups (p=0.0009) (Fig. 4B and Supplementary Fig. 4D). In agreement with these findings, the ipsilateral hippocampus of XYf and XXm animals showed reduced neuronal activity compared to the contralateral hippocampus (Paired t-test, p=0.0113 and p=0.02, respectively) (Fig. 4C, 4D and Supplementary Fig. 4E). These results align with those obtained using WT mice. Altogether, these results demonstrate that the presence of ovaries is not sufficient to protect memory-associated neuronal circuits, but that a XX chromosome complement is also necessary.

**Fig. 4:**
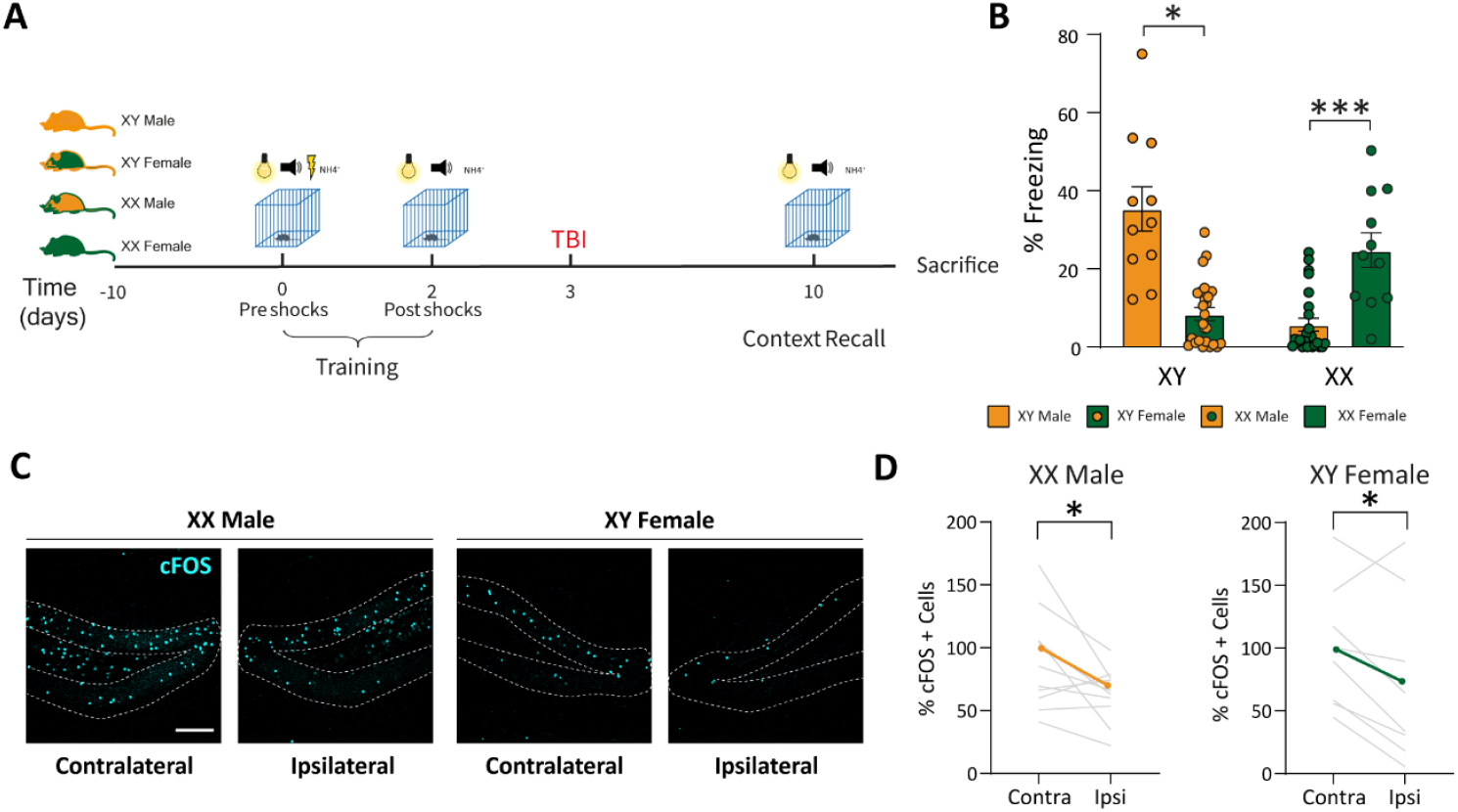
Sex-chromosomes counts to sex-differences in memory and hippocampal neuron activity after TBI. (**A**) Time course of the experimental procedure, like described in Figure 1A, except for castration and tibolone treatment. (**B**) Quantification of the freezing response in FCG animals (% of time). (**C**) Representative images of neuronal activation in the hippocampus by cFos immunostaining (cyan). (**D**) Quantification of the cFos+ cells in the contralateral and ipsilateral hippocampus. Data in B are presented as mean ± SEM; two-way ANOVA followed by Fisher’s multiple comparisons post hoc test, n ≥ 19 animals per group. Data in D are presented as individual values; paired t-test. n = Individual values for each histological section (equivalent histological sections from a minimum of 4 animals). *p < 0.05; **p < 0.01. Scale bar 200µm.

Next, we investigated astrogliosis in the perilesional cortex in FCG mice. GFP mRNA expression analysed by Two-way ANOVA showed an interaction effect between chromosomes and gonads (F=4.902, p=0.0408). XYf and XXm animals presented significantly reduced levels of GFAP mRNA expression compared to XYm (Unpaired t tests, p= 0.0325 for XYf and p=0.0364 for XXm) (Fig. 5A and 5B). Analysis of S100B expression showed similar results (Fig. 5C and 5D). Two-way ANOVA, F(interaction)=10.62, p=0.0044; F(sex)=5.259, p=0.0341; F(gonads)=12.64, p=0.0023), with XYf and XXm animals showing significantly reduced levels compared to XYm (Unpaired t test, p=0.0035 for XYf and p=0.0087 for XXm). We also analyzed Homer1 fluorescence intensity in astrocytes near the lesion by IHC in FCG model mice (Supplementary Fig. 5). As shown in Fig. 5E and 5F, animals with XX chromosomal complement and/or ovaries showed higher levels of Homer1 compared to XYm (Two-way ANOVA, followed by Fisher’s multiple comparisons test, F(interaction)=5.803, p=0.0347; F(sex)=8.41, p=0.0144; p= 0.0337 for XYf vs. XYm.

**Fig. 5:**
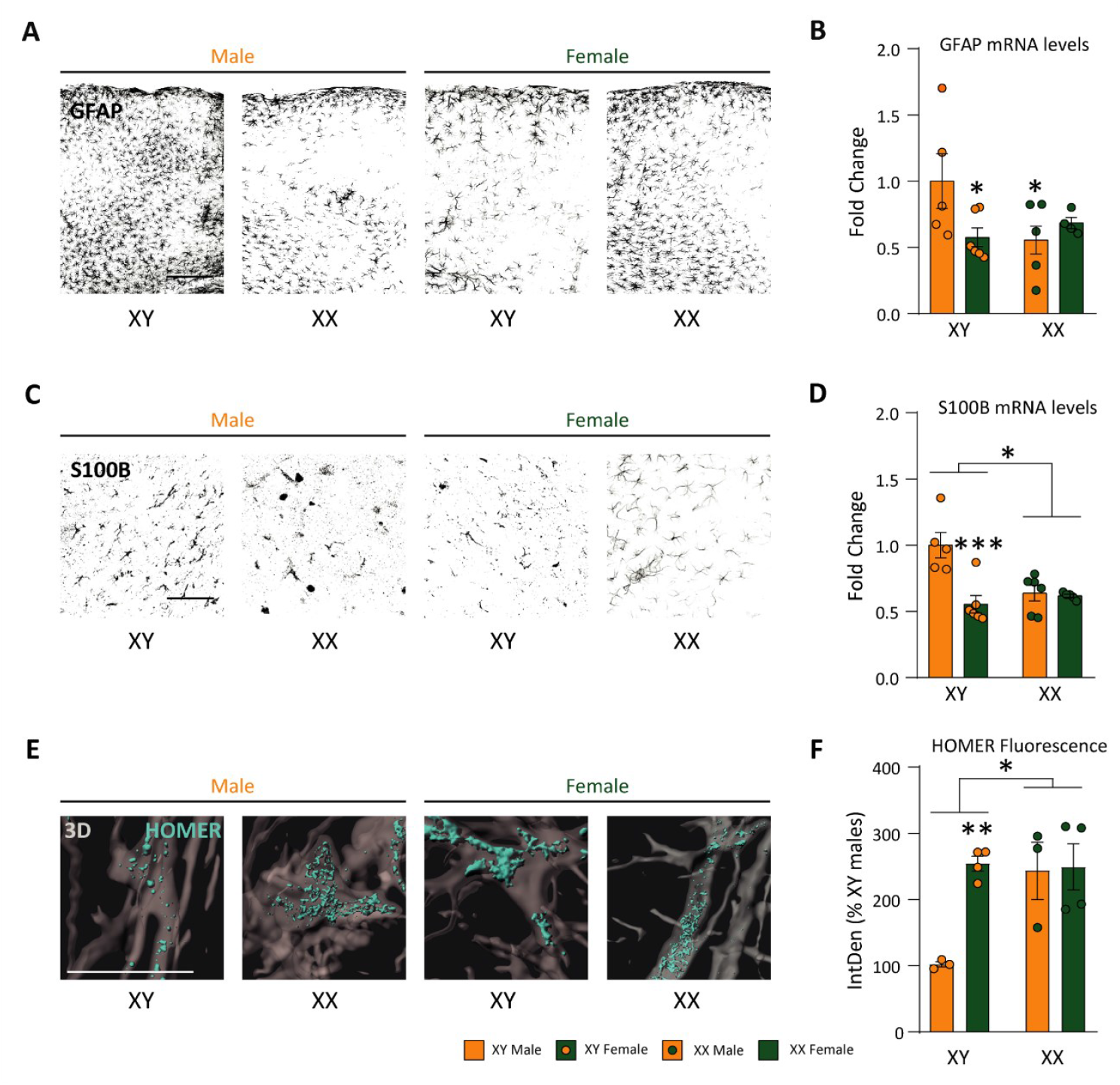
The presence of XX chromosomes or ovarian hormones ameliorates the astrogliosis produced by TBI. (**A, C**) Representative confocal images of GFAP (black) and S100B (black) IHC signal in the perilesional cortex. (**B, D**) Quantification of GFAP and S100B mRNA levels in the tissue lysates, measured by qRT-PCR. (**E**) IMARIS 3D reconstructions of Homer1 (cyan) expression in GFAP+ astrocytes (gray). (**F**) Quantification of the Homer integrated density (IntDen) inside the astrocytes. Data (percentage of XY male group) are presented as the mean ± SEM; two-way ANOVA followed by Fisher’s multiple comparisons post hoc test. n ≥ 3 animals per group. *p < 0.05; **p < 0.01; ***p < 0.001. Scale bars: 250µm in (A), 100µm in (C) and 15µm in (E).

In summary, the results obtained here prove that the presence of either XX chromosomes or ovaries contributes to partially reduce the astrocytic reactivity. Nevertheless, the combination of both XX chromosomes and ovaries is necessary to prevent reduction in neuronal activity in the hippocampus and subsequent memory loss.

### XYf and XXm astrocytes show defective myelin uptake and reduced levels of the phagocytic receptor Megf10

Finally, we assessed the impact of sex chromosomes on astrocytic phagocytosis of myelin (Supplementary Fig. 6A). As seen in Fig 6A and 6B, all groups presented significantly lower levels of myelin uptake compared to XXf (Two-way ANOVA, F(interaction)=14.94, p=0.0014; Unpaired t tests: p=0.0152 for XYm, p=0.0004 for XYf and p=0.0019 for XXm vs. XXf). These myelin engulfment levels were correlated with Megf10 expression (Fig. 6C and 6D, and Supplementary Fig. 6B). Two-way ANOVA evidenced a significant interaction between sex and gonads, F(interaction)=30.01, p<0,0001. Unpaired t tests showed a significant reduction of protein expression when animals had gonads that do not match with their sex chromosomes (p=0.0009, XYm vs. XYf, and p=0.0016 XXm vs XXf). Collectively, our data indicate that even though XX chromosomes or ovaries alone can partially reduce gliosis, unless combined, they fail to induce significant Megf10 receptor expression in astrocytes. This impairs phagocytosis of myelin debris, potentially creating a more adverse environment for hippocampal neurons, which can affect their activation and impair memory retention.

**Fig. 6:**
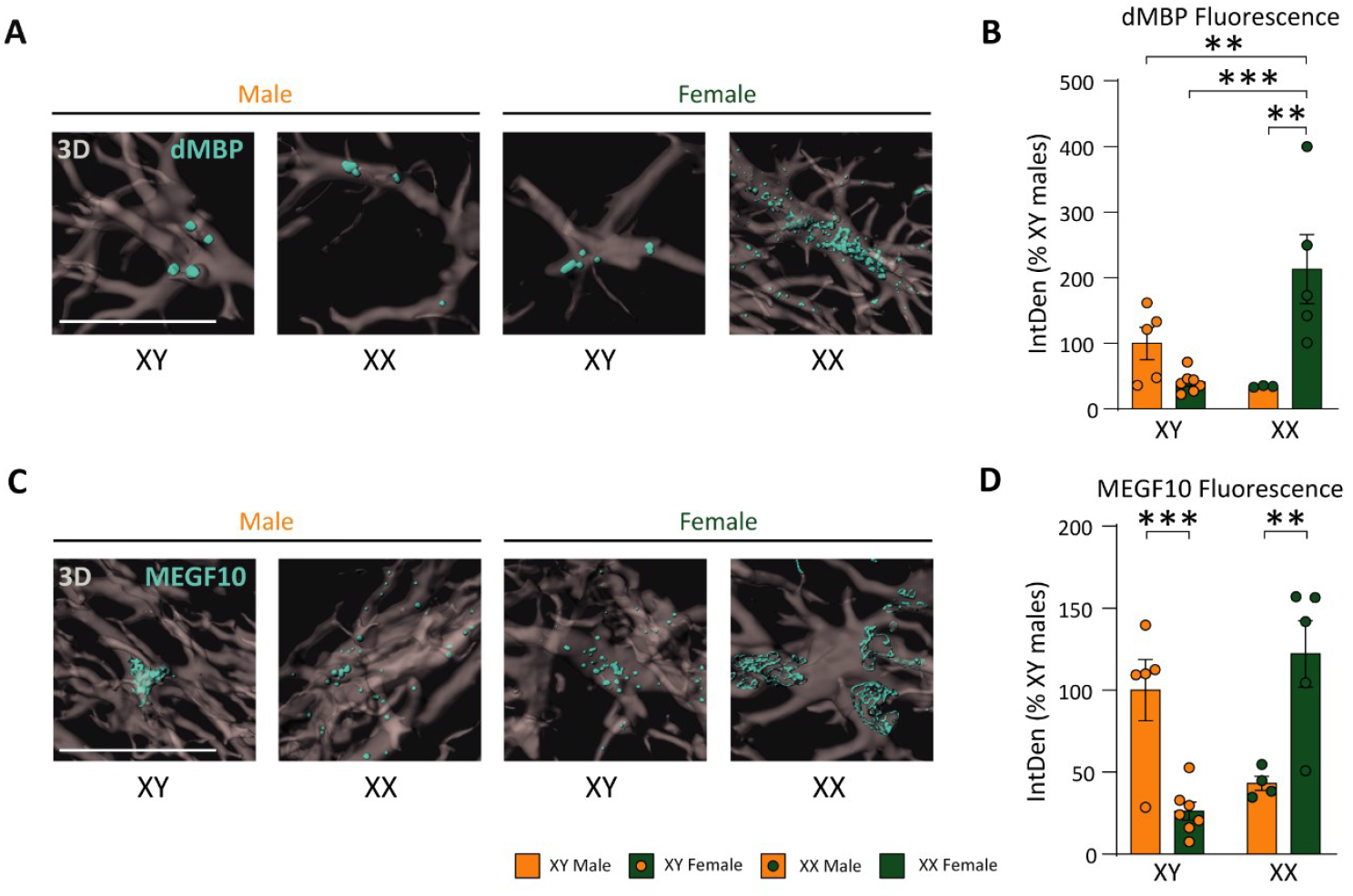
Myelin uptake and phagocytic receptor (Megf10) expression are reduced in XYf and XXm. (**A, C**) IMARIS 3D reconstructions of dMBP and Megf10 (cyan) within the astrocytes (gray). (**B, D**) Quantification of engulfed myelin and Megf10 protein expression was evaluated by measuring dMBP and Megf10 fluorescence intensity within GFAP+ astrocytes. Data are presented as the mean ± SEM; two-way ANOVA followed by Fisher’s multiple comparisons post hoc test. n ≥ 3 animals per group. *p < 0.05; **p < 0.01; ***p < 0.001. Scale bars 15µm.

### Astrocyte transplantation confirms sex-specific intrinsic and extrinsic regulation of astrocyte response in TBI

Male and female brains develop under distinct hormonal timelines, with an early postnatal androgen surge in males and a late estrogen surge in adult young females [29]. Early postnatal exposure to male hormones is thought to imprint transcriptional programs in microglia that predispose them to a pro-inflammatory state. To interrogate whether male and female astrocytes display different intrinsic abilities to cope with an insult, irrespective of other brain cells we performed sex-matched and cross-sex astrocyte transplantation. Primary astrocytes obtained from Aldh1l1-EGFP mice were injected into the region of the cerebral cortex designated for TBI (Fig. 7A). We first validated the experimental approach by confirming the survival and correct localization of transplanted astrocytes in the host brain i.e., at the same distance to injury as the endogenous astrocytes analyzed in previous experiments. Transplanted astrocytes expressed the astrocytic marker GFAP and extended hypertrophic processes, suggesting active engagement with the host cells and the extracellular environment, indicative of a successful integration one-week post-transplantation (wpt) (Fig. 7B).

**Fig. 7:**
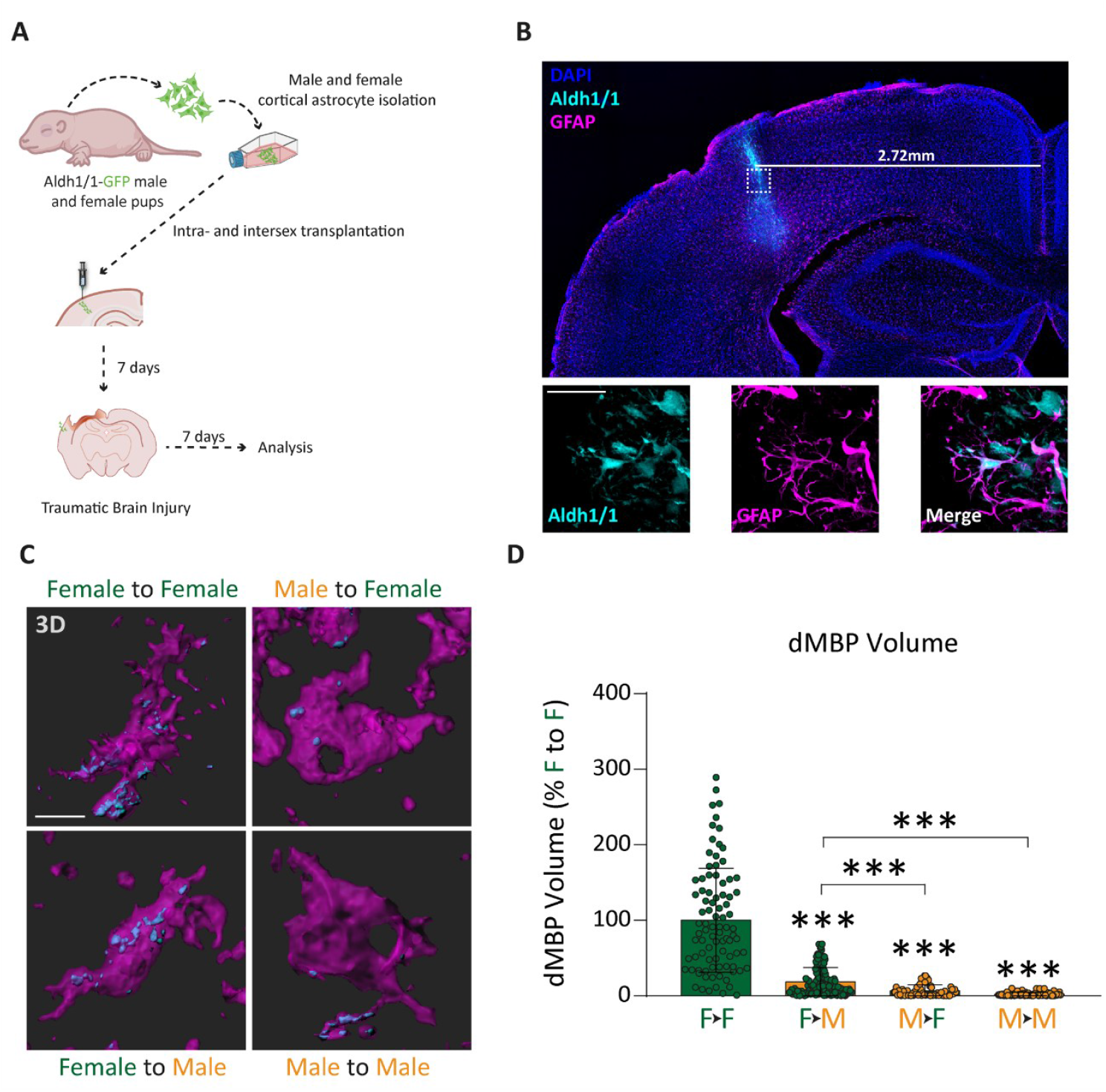
Transplanted female astrocytes exhibit enhanced phagocytosis of myelin debris. (**A**) Scheme of the experimental protocol. Fluorescent astrocytes from Aldh1/1-GFP male and female pups were isolated and transplanted into the cerebral cortex of adult male and female wildtype animals. Seven days later, TBI was performed near the transplant site. (**B**) Representative confocal tile-scan showing the transplanted astrocytes (cyan) in the target area. The zoom-in images (bottom) confirm that the transplanted cells are astrocytes, as they express the astrocytic marker GFAP (magenta). (**C**) IMARIS 3D reconstructions of dMBP (cyan) within astrocytes (magenta). (**D**) Quantification of myelin uptake was evaluated by measuring dMBP volume within GFAP+ astrocytes. Data are presented as mean ± SEM; nonparametric one-way ANOVA followed Dunn’s multiple comparisons post hoc test for each sex. n = 65-126 cells of at least 3 animals per group. ***p < 0.001. Scale bars: 50µm in (B) down, 15µm in (C).

To determine the contribution of brain environmental signals to previously observed sex differences in myelin phagocytosis, we injected the cells one week before the injury into the brains of same-sex and opposite-sex mice. One week after injury, we assessed the capacity of transplanted astrocytes to phagocyte myelin. As previously, 3D reconstruction of GFAP and dMBP fluorescence signals from confocal images enabled quantification of dMBP amount contained within the transplanted astrocytes (Fig. 7C). Results show that female astrocytes introduced into female brains exhibited significantly higher phagocytic activity compared to all other experimental groups (Unpaired t tests, p<0.0001 for all comparisons) (Fig. 7D). Moreover, female astrocytes transplanted into male brains phagocytosed significantly more myelin debris than male astrocytes regardless of the host environment, confirming that both sex-specific cell-autonomous programs and non-cell autonomous factors regulate astrocyte phagocytic activity.

## Discussion

Biological sex has been proposed as an important modifier of TBI outcome, with females presenting better recovery. Here, we present novel evidence that both gonadal hormones and sex chromosome complement are critical determinants of the cellular and cognitive consequences of a cortical injury in mice. By combining two complementary experimental strategies: gonadectomy in wild-type (WT) animals and the use of Four Core Genotypes (FCG) mice, we were able to disentangle the contributions of the hormonal environment and the chromosomal endowment to TBI-induced contextual memory deficits and associated cellular responses (neuronal and astrocytic) in the hippocampus and cortex. Furthermore, sex-matched and cross-sex astrocyte transplantation experiments demonstrated that both intrinsic sex-specific astrocyte properties and extrinsic sex-specific signals play a role in debris clearance by astrocytes, a process that facilitates neuronal survival and function. Collectively, our findings position astrocytes as central players in sex differences following injury, a role that, unlike microglia, has yet remained largely unexplored.

Our findings first demonstrate that, following TBI, freezing behavior diverges in a sex-dependent manner. Males exhibited a marked decrease in freezing, whereas females maintained performance, indicating that males are more vulnerable to TBI-induced memory impairment, an observation corroborated by previous evidence [30, 31]. Gonadectomy and tibolone treatment revealed that the female advantage requires circulating ovarian hormones: ovariectomy reduced freezing times in females after TBI, matching the male phenotype, while tibolone supplementation restored performance. Males, however, showed no alterations with neither gonadectomy nor tibolone, suggesting sex differences in steroid hormone receptors: expression levels, distribution throughout the brain and/or in their downstream signaling pathways. In line with this view, previous studies described significant sex bias in ERα and ERβ expression patterns [32, 33] and in gene transcription downstream activation of ERα [34] and ERβ [35]. Experiments in FCG mice further supported this view, as the presence of ovaries in XY animals did not improve scores. It remains elusive, however, whether the genetic advantage is mediated by the complement of genes encoded on the sex chromosome X, that escape X chromosome inactivation and contribute to sexually divergent features, or it results from permanent structural and functional divergences in gene expression and neural circuits established during brain sexualization, a process governed by both gonadal hormones and sex chromosomes. Notably, literature suggests that developmentally established sex differences can impact vulnerability to perturbations or disease [36-38].

Secondly, our behavioral results were paralleled by levels of hippocampal neuronal activity as measured by cFos immunoreactivity. Females with functional ovarian hormone signaling exhibited higher cFos+ neuronal density after injury compared to males or ovariectomized females. These findings suggest that ovarian hormones preserve neuronal activity and fear memory, and agree with previous reports [39]. Estrogen, has been strongly implicated in neuroprotection after TBI through multiple mechanisms and notably, neuroprotection is more robust in females [2, 40]. This is thought to be because of 1) higher basal ERs expression in key cognitive regions such as the hippocampus, 2) greater estradiol availability from ovaries in females versus the aromatization-dependent supply in males and 3) differences in ER downstream signaling. Importantly, estradiol-mediated neuroprotection was initially thought to be mediated directly through neuronal ERs, however, compelling data now also implicate astrocytes and microglia in ER-mediated neuroprotection by controlling neuroinflammation [41-43].

While a direct role of estrogen signaling in astrocytes in promoting neuronal survival is established, their influence on gliosis and debris clearance is less explored, processes that significantly shape the injury microenvironment and the resilience of neuronal networks. Herein, we evaluated astrocyte reactivity by GFAP, S100B, and Homer1 expression. GFAP levels increased after TBI in both sexes, consistent with our earlier observations in stab wound injury, where no sex differences in GFAP+ cell number were found [27]. Nevertheless, here we show that females displayed less astrogliosis than males as revealed by lower S100B expression, an indicator of injury severity and neuroinflammation [44, 45], that was reduced by approximately 40% in injured females with ovaries compared to injured males and ovariectomized females. Tibolone treatment reduced astrogliosis, pointing to a role of estrogen in attenuating astrocytic reactivity [4, 46]. XXm animals also showed diminished GFAP and S100B expression, compared to XYm, suggesting that XX chromosomes can partially limit gliosis independently of hormonal status. Homer1, associated with a neuroprotective astrocytic phenotype capable of limiting excitotoxic glutamate signaling [28, 47, 48] was higher in female astrocytes and reduced after gonadectomy, with tibolone restoring injury levels. Homer1 upregulation was also observed in XXm and XYf mice pointing to both chromosome- and hormone-linked protection. Together, these data indicate that both ovarian hormones and XX chromosomes contribute to reduce astrogliosis and promote neuroprotective astrocyte phenotypes, although their effects are not strictly additive.

Rapid removal of cellular debris is crucial for restoring tissue homeostasis and for preventing excessive inflammatory responses that can propagate damage to surrounding tissue. While microglia are well-known mediators of debris clearance, accumulating evidence shows that reactive astrocytes can also engulf a variety of cellular components, including synapses, axons, and apoptotic cells through phagocytic receptors ABCA1, Megf10 and Mertk [49-52]. Notably, astrocyte-mediated clearance of myelin debris was also sex-dependent. Engulfment correlated with Megf10 expression and was more pronounced in females, diminished after ovariectomy, with tibolone partially restoring phagocytosis. Although Megf10 receptor has not previously linked to myelin phagocytosis, myelin debris are rich in complex lipids, and phosphatidylserine was suggested as the “eat me” signal of myelin debris [11]. In FCG mice, XXf exhibited the highest phagocytic activity, indicating that both XX chromosomes and ovarian hormones synergize to optimize debris clearance. Mismatch between hormonal environment and chromosome complement (XYf and XXm) reduced phagocytosis, suggesting that coordinated hormone-gene interactions are necessary for maximal astrocytic clearance. These findings mirror the performance in the fear conditioning test and cFos immunoreactivity, supporting the idea that efficient debris removal contributes to neuronal and cognitive protection after TBI.

To unequivocally determine whether sex dimorphism in astrocyte-mediated myelin clearance arises from intrinsic differences between male and female astrocytes, rather than from sex differences in other cells that signal to astrocytes (e.g. microglia) we transplanted astrocytes between sexes. Female-derived astrocytes retained superior phagocytic capacity regardless of the host sex, indicating that sex-linked gene expression, likely X chromosome dosage effects or permanent effects of brain sexualization, confers a baseline advantage in debris clearance. Yet, female brains supported the most robust phagocytosis, indicating that extracellular signals further modulate this process (such as hormonal supply or molecular signals derived from other cell types). These data corroborate our findings obtained in WT and FCG mice, underscoring the divergence of cell-intrinsic myelin clearance properties between female and male astrocytes, which are further regulated by external signals such as circulating hormones.

Overall, our results demonstrate that protection from TBI-induced cognitive impairment in females arises from the combined influence of ovarian hormones and XX chromosome complement. This synergy limits astrocytic reactivity, enhances myelin debris clearance, and preserves hippocampal neuronal activity and memory. Neither factor alone fully replicates the female advantage. Importantly, tibolone restored several female-specific protective features but was ineffective in males and carries potential cancer risks. Short-term therapeutic windows of tibolone treatment may mitigate these risks while offering neuroprotective benefits. On the other hand, our work points at other promising therapeutic approaches namely targeting astrocytic estrogen receptor signaling, Homer1 levels or Megf10-mediated phagocytosis.

## Conclusion

This study provides the first evidence that both gonadal hormones and sex chromosome complement jointly determine astrocyte response to injury and associated cognitive outcomes after TBI. Considering both dimensions of biological sex, dynamic hormone changes over the lifespan and integrating glial interactions will be essential for accurate modeling of brain injury and the development of effective, personalized interventions.

## Supplementary Information

This article has an accompanying supplementary file: Supplementary Material 1.

## Declarations

### Ethics approval and consent to participate

All the procedures applied to the animals used in this study followed the European Parliament and Council Directive (2010/63/EU) and the Spanish regulation (R.D. 53/2013 and Ley 6/2013, 11th June) on the protection of animals for experimental use and were approved by our institutional animal use and care committee (Comité de Ética de Experimentación Animal del Instituto Cajal) and by the Consejería del Medio Ambiente y Territorio (Comunidad de Madrid, PROEX 059.5/21).

### Consent for publication

Not applicable

### Availability of data and materials

All data generated or analyzed during this study are included in this published article (and its supplementary information files)

### Authors’ contributions

DPB, SG and MAA conceived and designed the study. DPB, CPL, NCA and IA generated the data. DPB, DG, FPUS, SG and MAA processed and analyzed the data. DPB, SG and MAA wrote the manuscript, with input from all authors. All authors read and approved the final version of the manuscript.

## Acknowledgements

We thank all the facilities at the VBC without which this work would not have been possible, in particular the animal house and bio-optics. We are particularly grateful to Anna Smolka for technical assistance in this work.

