## Supplementary Figures for "Improved myelin clearance and cognitive outcomes after TBI in female mice are mediated by ovarian steroids and sex chromosomes"

**Supplementary Fig. 1**

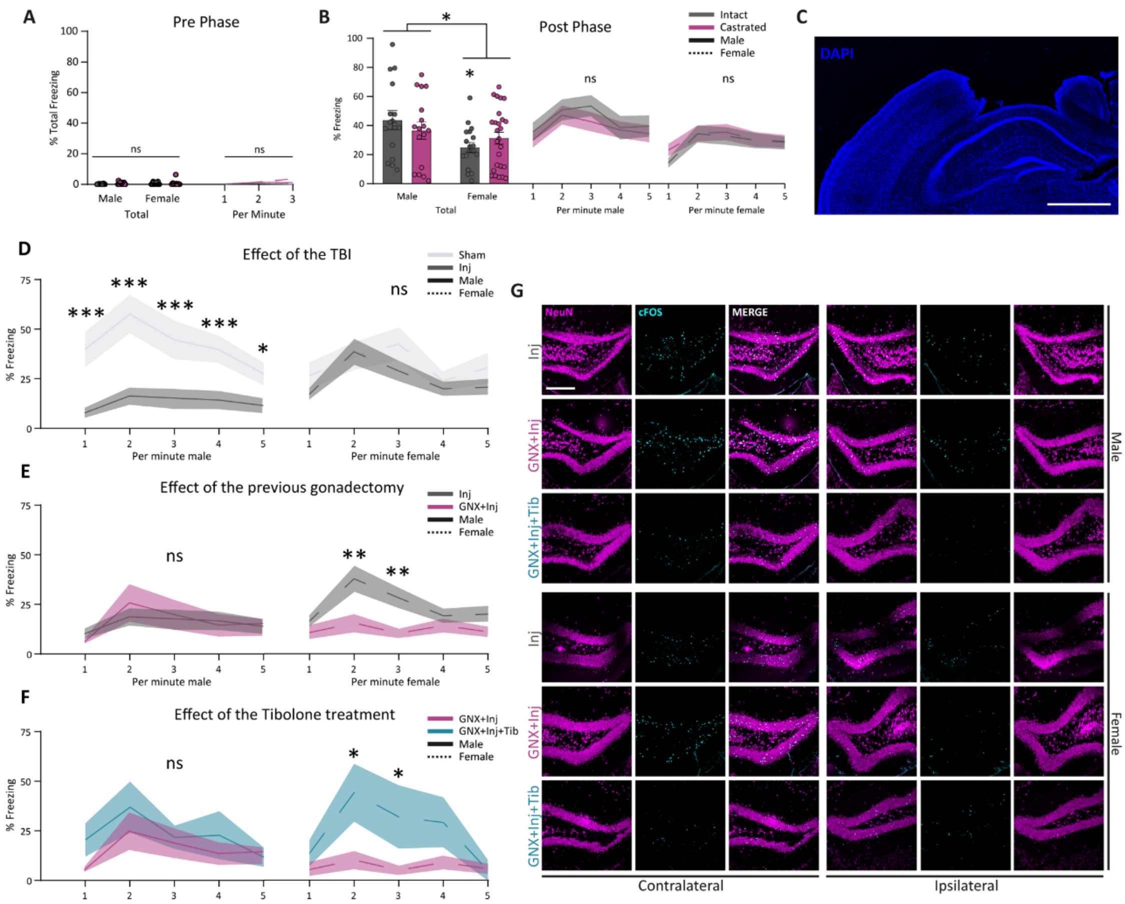

**Fig. S1:** (**A**) Percentage of freezing time for the experimental groups before receiving the aversive stimulus (Pre-phase). Note the absence of freezing behavior. (**B**) Percentage of total (left) and minute-detailed (right) freezing time of the animals during the recall phase before the TBI. Castration has no effect on the memory of the animals before being submitted to the injury. (**C**) Representative confocal tile-scan showing the cortical injury above medial hippocampus (DAPI in blue). (**D, E, F**) Minute-detailed freezing time of animals submitted to an injury (Sham vs. Inj), castration and injury ((Inj vs. GNX+Inj) and tibolone treatment upon castration and injury (GNX+Inj vs. Tib+GNX+Inj), respectively. (**G**) Confocal images of the hippocampus showing the activated neurons. Neuronal marker NeuN (magenta) and cFos marker of activation (cyan) were used. Data are presented as mean ± SEM; two-way ANOVA followed by Fisher´s multiple comparisons post hoc test. n ≥ 8 animals per group. *p < 0.05; **p < 0.01; ***p < 0.001. Scale bars: 1mm in (A), and 200µm in (G).

**Supplementary Fig. 2**

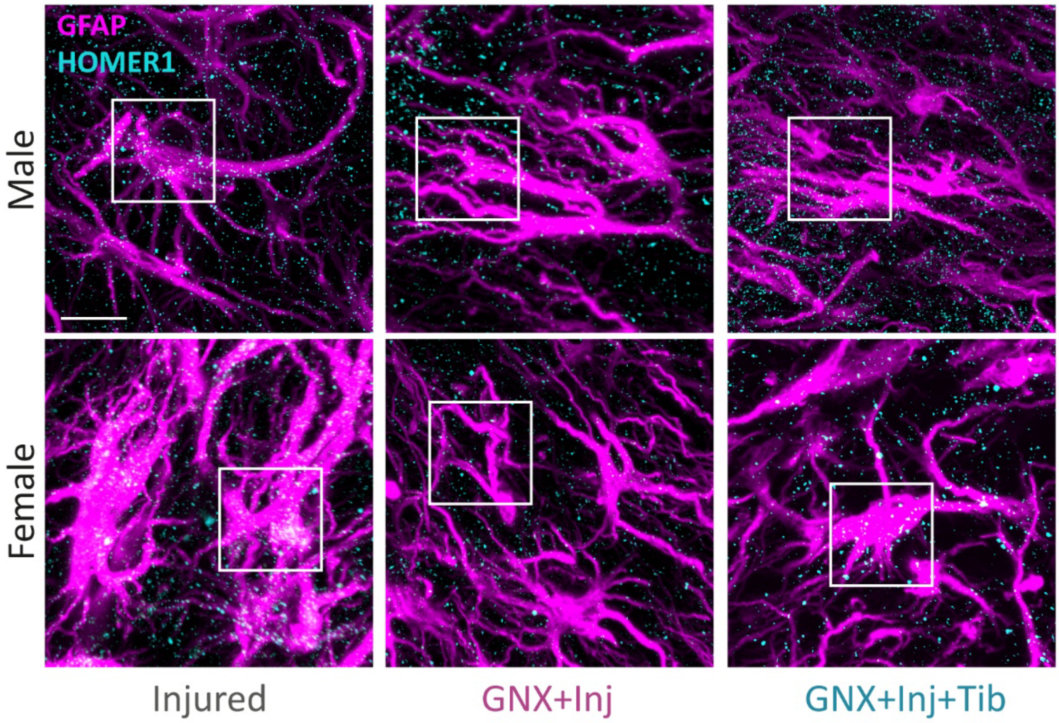


**Fig S2.** Confocal images show Homer1 (cyan) expression by astrocytes stained with astrocytic marker GFAP (magenta). Scale bar: 15μm.

**Supplementary Fig. 3**

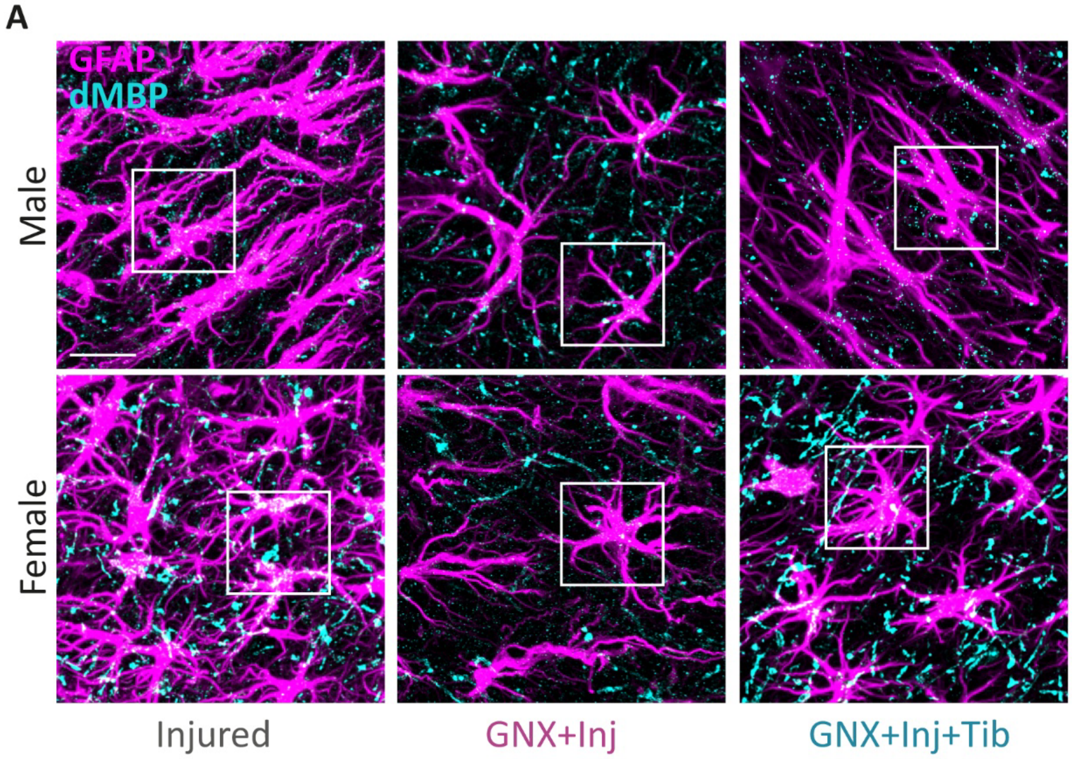


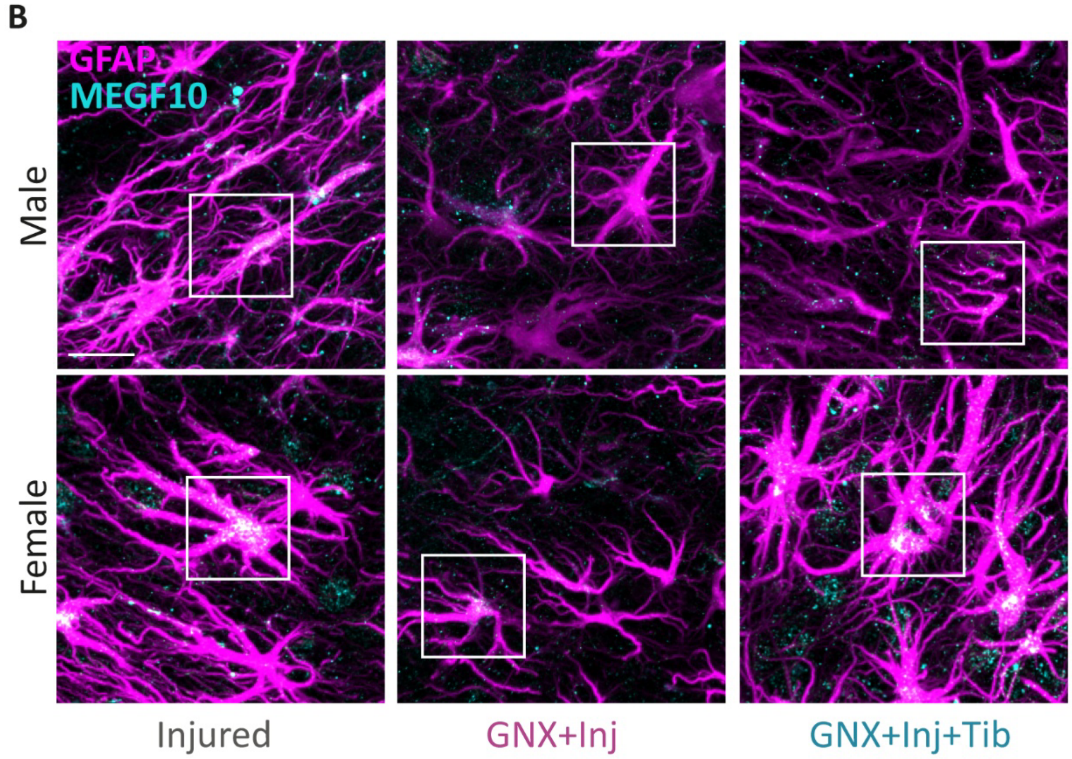


**Fig. S3: (A)** Confocal images show dMBP (cyan) signal within astrocytes stained with astrocytic marker GFAP (magenta) indicating phagocyted myelin debris. **(B)**

Confocal images show Megf10 phagocytic receptor (cyan) expressed by astrocytes stained with astrocytic marker GFAP (magenta). Scale bars: 15µm.

**Supplementary Fig. 4**

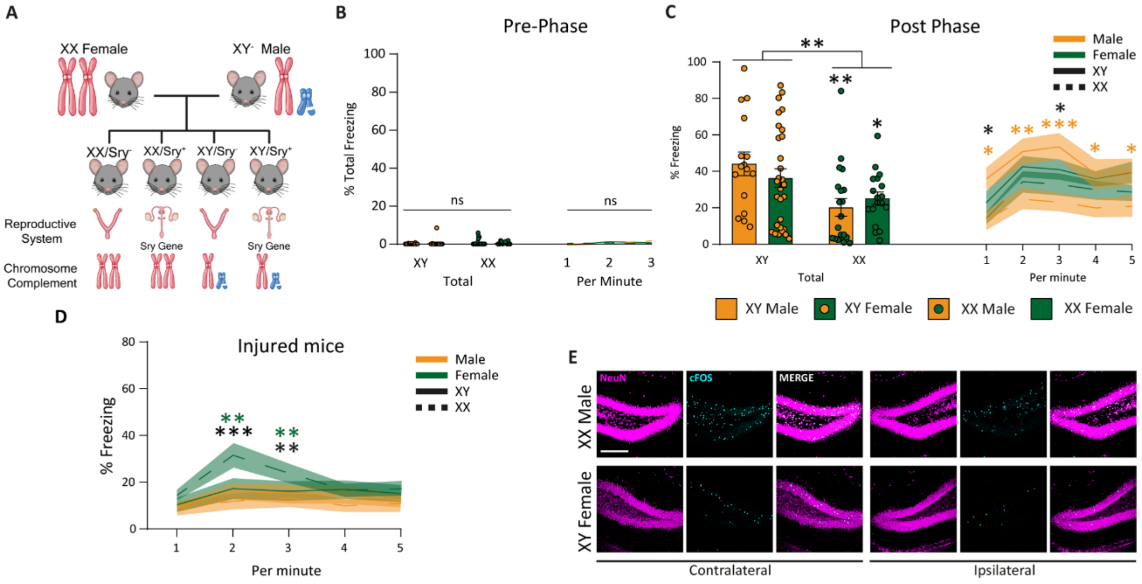


**Fig. S4. (A)** Schematic of the FCG mice. By crossing a WT female with a XY male containing the *Sry* in chromosome 3, the progeny results in XY animals with ovaries or XX animals with testicles in addition to XX with ovaries and XY with testicles. **(B)** Percentage of freezing time (total) of the FCG mice before receiving the aversive stimulus (Pre-phase). Note the absence of freezing behavior. **(C)** Total (left) and minute-detailed (right) freezing time of the animals during the recall phase before the TBI. Freezing levels of XX animals are significantly lower. **(D**) Minute-detailed freezing time of FCG animals submitted to an injury. **(E)** Confocal images of the hippocampus showing the activated neurons. Neuronal marker NeuN (magenta) and cFos marker of activation (cyan) were used. Data are presented as the mean ± SEM; two-way ANOVA followed by Fisher´s multiple comparisons post hoc test. n ≥ 19 animals per group. *p < 0.05; **p < 0.01; ***p < 0.001. Yellow and green asterisks compare XYm vs XXm, and XYf vs XXf, respectively. Scale bar: 200µm.

**Supplementary Fig. 5**

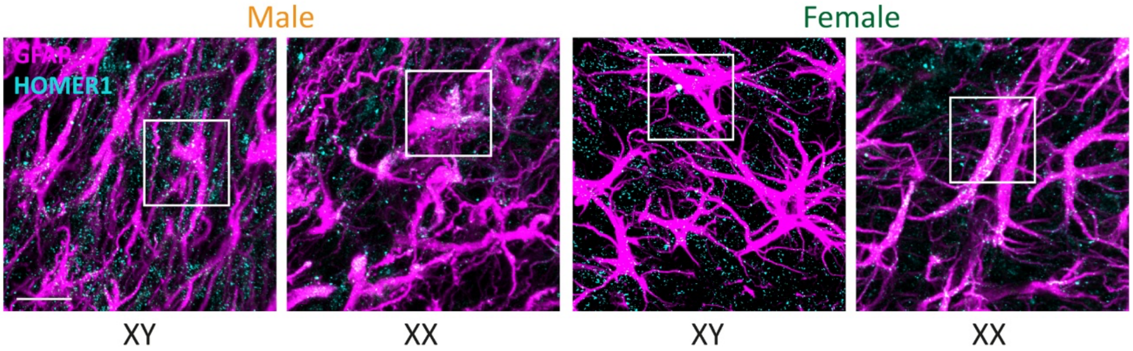


**Fig. S5:** Confocal images of Homer1 (cyan) expressed by astrocytes stained with astrocytic marker GFAP (magenta) in FCG mice. Scale bar: 15μm.

**Supplementary Fig. 6**

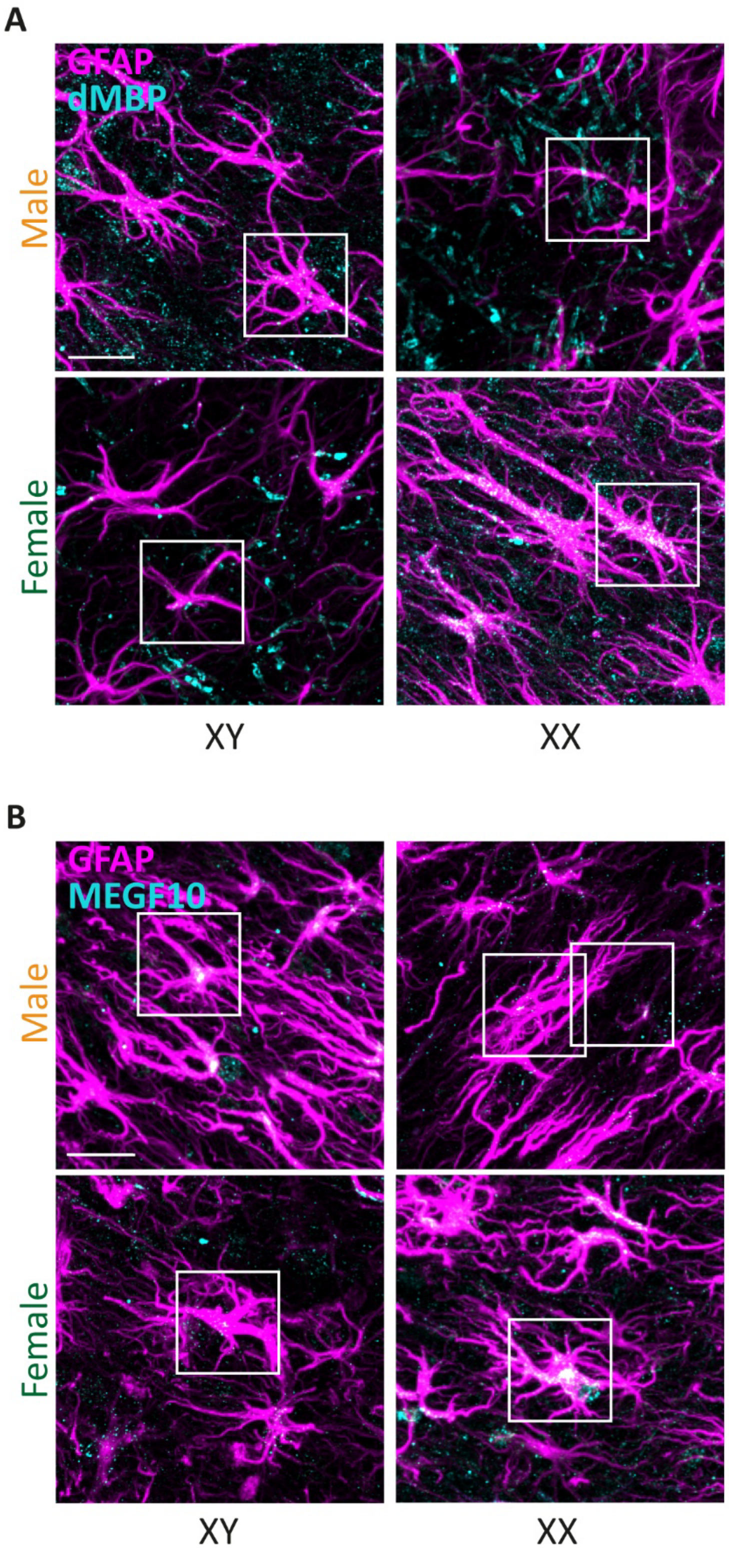


**Fig. S6: (A)** Confocal images show dMBP (cyan) within astrocytes stained with astrocytic marker GFAP (magenta) in FCG mice, indicative of myelin phagocytosis. **(B)** Confocal images show Megf10 phagocytic receptor (cyan)

expressed by astrocytes stained with astrocytic marker GFAP (magenta) in FCG mice. Scale bars: 15µm.

**Supplementary Fig. 7**

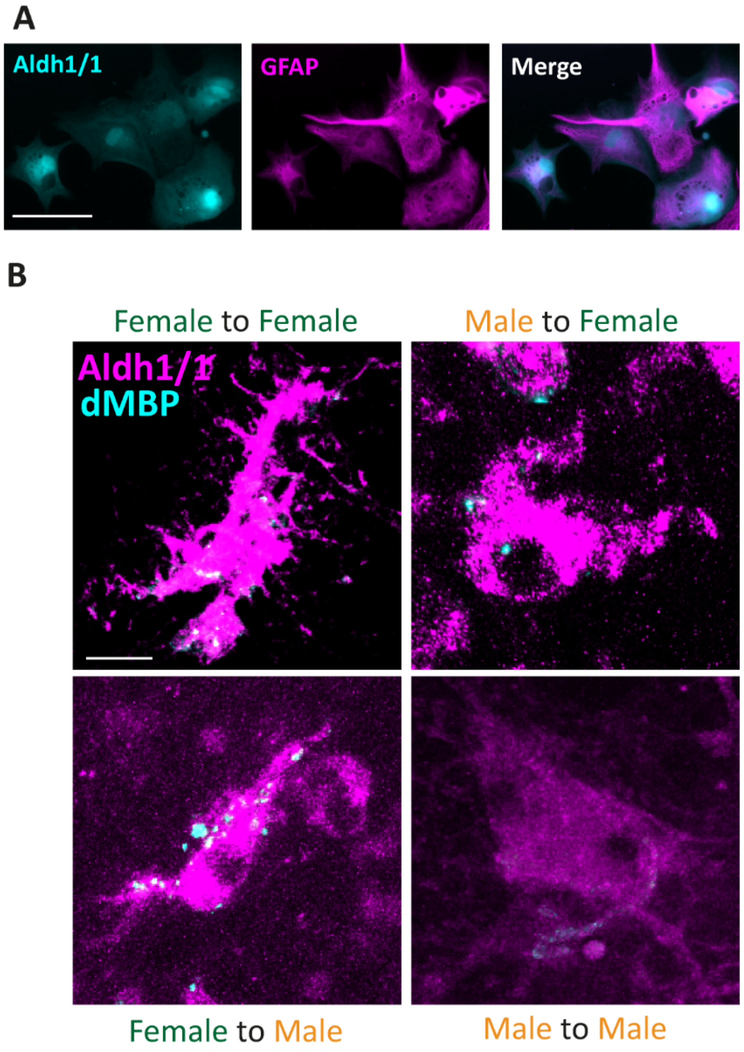


**Fig. S7: (A)** Primary cultures of cortical astrocytes obtained from Aldh1/1-GFP mice. The colocalization of the endogenous fluorescent reporter GFP (cyan) and the astrocytic marker GFAP (magenta) confirm that the isolated cells are astrocytes. **(B)** Confocal images show intra- and cross-sex transplanted astrocytes expressing the endogenous reporter Aldh1/1 (magenta) and phagocyting dMBP (cyan). Scale bars: 15µm in A and 5µm in B.
